# Deep generative model for protein subcellular localization prediction

**DOI:** 10.1101/2024.10.29.620765

**Authors:** Guo-Hua Yuan, Jinzhe Li, Zejun Yang, Yao-Qi Chen, Zhonghang Yuan, Tao Chen, Wanli Ouyang, Nanqing Dong, Li Yang

## Abstract

Protein sequence determines not only its structure but also its subcellular localization. Although a series of artificial intelligence models have been reported to predict protein subcellular localization, most of them provide only textual outputs. Here, we present deepGPS, a <u>deep</u> <u>g</u>enerative model for <u>p</u>rotein <u>s</u>ubcellular localization prediction. After trained with both protein primary sequences and protein subcellular localization fluorescence images, deepGPS shows the ability to predict cytoplasmic and nuclear localizations by reporting both textual labels and generative images as outputs. In addition, deepGPS shows potential to be further extended for other types of subcellular localization prediction, even with limited input data volumes for training. Finally, an openGPS website (https://bits.fudan.edu.cn/opengps) is constructed to provide a public and convenient platform for protein subcellular localization prediction with the scientific community.

## Introduction

Proteins, as the ultimate effectors in the central dogma, perform essential functions in living cells and organisms, including but not limited to providing structural support, catalyzing biochemical reactions, defending against pathogens, transmitting signals and regulating gene expression ^1–4^. The precise subcellular localization of proteins within the cell determines their ability to interact with appropriate molecules for correct functioning, while their abnormal localization may link with uncontrolled roles in cell and in the progress of human diseases, such as Alzheimer’s disease and cancers ^5, 6^. Thus, studying protein subcellular localization is crucial for understanding their function in different biological and pathological contexts.

Generally, protein subcellular localization can be identified through different experimental methods, such as fluorescence tagging or Western blotting after fractionation ^7^. These types of protein subcellular localization information are collected and recorded in the public protein database like UniProt ^8^, usually with text labels, and shared in the community. With these protein subcellular localization data in hand, several computational methods based on machine learning and deep learning models have been also developed to predict protein subcellular localization. For example, as early as in 2001, Hua and Sun constructed a support vector machine (SVM) model to predict the subcellular localization of proteins from amino acid compositions ^9^. In addition, the machine learning model MultiLoc2 ^10^ predicts protein subcellular localization using amino acid composition, phylogeny and Gene Ontology terms based on the SVM framework. Furthermore, a deep learning model DeepLoc ^11^ employs a recurrent neural network and the attention mechanism to predict protein subcellular localization. Although these artificial intelligence models predict protein localization with reasonable performance, they only output textual format labels of subcellular localization information of given proteins (Extended Data Fig. 1a). Thus, this text-to-text prediction lacks a direct visual representation like experimentally generated protein subcellular localization images.

Recently, Cho *et al*. constructed the OpenCell database ^12^, which maps the subcellular localization of 1,311 proteins using high-throughput CRISPR-mediated genome editing and live-cell fluorescence imaging. Utilizing these extensive and systematic data of protein localization in both text and image formats from OpenCell, two independent self-supervised models^12, 13^ have been developed to perform image-to-image prediction. These models are designed to reconstruct the fluorescence images that closely resemble the true images, capturing intrinsic protein localization patterns through the features learnt by the models (Extended Data Fig. 1b) ^12, 13^. However, these models primarily focus on profiling and clustering subcellular localization based on existing fluorescence images^12, 13^. That is to say, these models focus on image analysis to identify hidden features, and thus could not be used to predict protein subcellular localization directly from the protein sequence. Therefore, a text-to-image prediction model of protein subcellular localization has been still desired.

In this study, we present deepGPS, a deep generative model for *de novo* protein subcellular localization prediction. Distinct to other previously reported AI models, deepGPS outputs both textual labels and artificial fluorescence images after prediction. After training with high quality fluorescence images from OpenCell database ^12^ together with their primary sequences and subcellular localization information of 1,311 proteins, deepGPS achieves highly reliable prediction outcomes for cytoplasmic and nuclear localized proteins with its text-to-image functionality, and also shows potential for other types of subcellular localization prediction even with limited input data volumes. Furthermore, we have constructed the openGPS website (https://bits.fudan.edu.cn/opengps), offering a public and convenient platform for protein subcellular localization prediction from the scientific community. We expect that our work will provide new insights into protein localization research, thus contributing to the understanding of protein function.

## Results

### An overview of deepGPS design

A deep generative model, deepGPS, was developed to predict protein subcellular localization. Different to other existing prediction models that input protein sequences and output a text label of localization^10, 11^, deepGPS incorporates protein subcellular localization image data from OpenCell database ^12^ together with protein primary sequences and subcellular localization information for model training. As a consequence, deepGPS outputs both text labels and artificial images for protein subcellular localization prediction (Fig. 1).

**Fig. 1:**
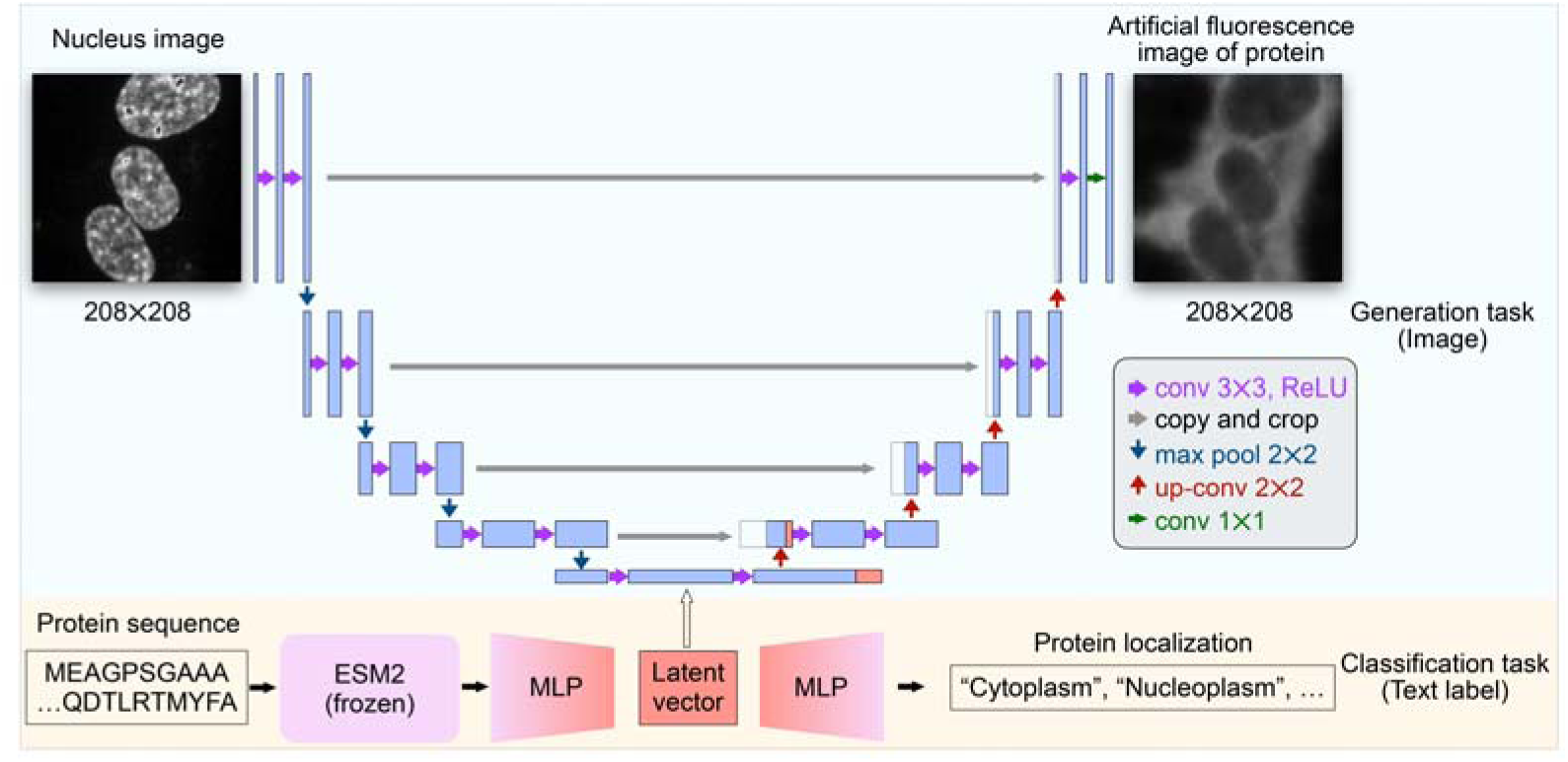
An overview design of deepGPS. A schematic diagram illustrating the architecture of deepGPS with the nucleus image and protein sequence as inputs. DeepGPS predicts protein subcellular localization, generating both a text label and an artificial fluorescence image as outputs.

Specifically, in the deepGPS model, the protein sequence was first encoded to a 1,280-dimensional vector by ESM-2 ^14^, a pre-trained large language model (LLM) of protein that has learned complex internal representations of protein sequences and achieved accurate predictions in protein structure prediction and protein-protein interaction ^15^. The encoded vector was then passed to a multilayer perceptron (MLP) with four layers and further output to the predicted text label of protein localization (Fig. 1, bottom). Meanwhile, each of 208 × 208-dimensional nuclei images from OpenCell database ^12^ was first encoded to a 64×208×208 dimension by a convolutional layer and then converted to a 1024×169-dimensional image latent vector through a series of down-sampling steps. This image latent vector was combined with the protein sequence latent vector from the MLP and further generated to a 208 × 208-dimensional protein localization image encompassing the given nuclei through up-sampling steps in the U-Net architecture (Fig. 1, top). With both the true text label and image of protein localization, we could then define a loss function by minimizing the difference between the true data and the predicted output to improve the performance of deepGPS (See “Methods” section).

### Construction of deepGPS with a comprehensive image dataset for protein subcellular localization

Since the performance of a model largely depends on the quality of the training dataset, we thus set to build a comprehensive image dataset of protein subcellular localization for deepGPS construction. Firs of all, we selected 1,301 out of 1,311 endogenously labeled proteins from the OpenCell database ^12^ as the whole protein dataset, which have complete predicted structures available from AlphaFold2 database^16^ (See “Methods” section). With these 1,301 proteins, we collected 6,239 paired images of protein fluorescence and corresponding nuclear fiducial marker from the OpenCell database ^12^. Next, to expand the sample size of training data, each pair of original images that contain multiple cells with clear fluorescence signals was processed into multiple pairs of cropped images through a series of procedures, including image normalization, nucleus segmentation, image denoising, and image cropping (Extended Data Fig. 2a, See “Methods” section). Briefly, after normalizing the raw intensities of the original images, the normalized images were segmented by nuclei using StarDist ^17^. Multiple regions of dimension 208 × 208 pixels were computationally chosen so that at least one nucleus is present and centered, resulting in a total of 62,108 cropped images of nuclei and paired protein localizations (Fig. 2a). Finally, we split this comprehensive image dataset into a 4:1 ratio using stratified sampling to ensure class uniformity in both the training and test sets (See “Methods” section). These data were used for subsequent model training and evaluation.

**Fig. 2:**
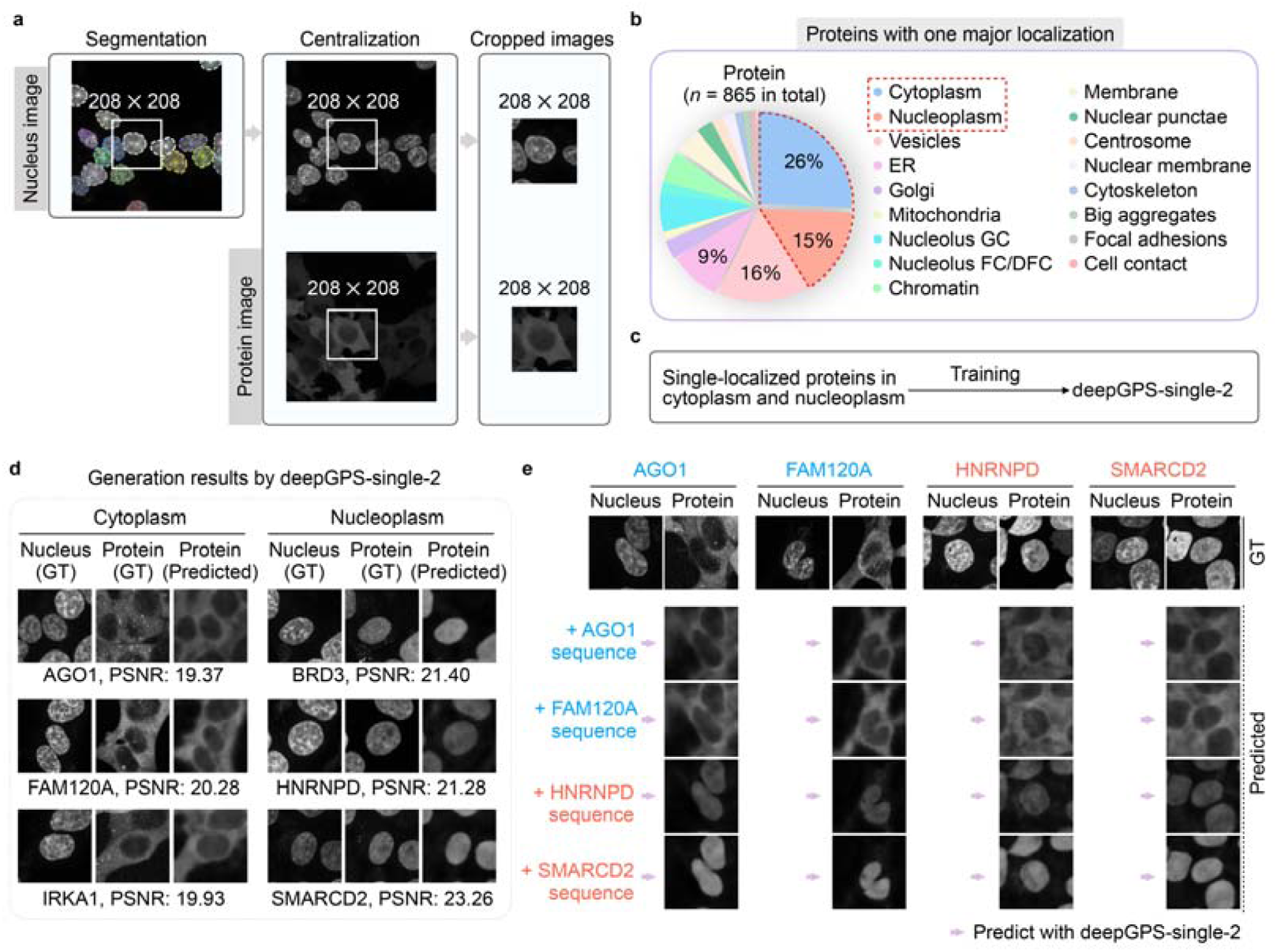
Construction and evaluation for deepGPS-single-2. **a**, Specific instance of image processing from segmented images to cropped images. **b**, Distribution of proteins with only one major localization. **c**, Strategy for training deepGPS-single-2. **d**, Six instances of generation results by deepGPS-single-2. GT, ground truth; PSNR, peak signal-to-noise ratio. **e**, A cross-assay using the ground-truth nucleus image as a nuclear fiducial marker and inputting different protein sequences to deepGPS-single-2. AGO1 and FAM120A are cytoplasmic proteins with color blue, while HNRNPD and SMARCD2 are nuclear proteins with color red. Of note, the proteins corresponding to images generated by deepGPS in panels **d** and **e** were all from test set, with sequences that were not used during deepGPS training.

Of note, most of the examined proteins (865 out of 1,301) exhibited only a single major subcellular localization, with similar ratios of images and cropped images (Extended Data Fig. 2b), which were used for subsequent model training. However, 41,261 cropped images of these 865 single-localized proteins were unevenly distributed across 17 organelles, primarily in four well-studied organelles, including cytoplasm (222 out of 865, ∼26%), nucleoplasm (132 out of 865, ∼15%), vesicles (140 out of 865, ∼16%) and endoplasmic reticulum (ER) (78 out of 865, ∼9%) (Fig. 2b and Extended Data Fig. 2c). In contrast, the remaining single-localized proteins (293 out of 865, about 34%) were sparsely distributed across other 13 sub-organelles.

Given that the uneven data distribution poses a challenge for achieving robust multi-class classification, we first tested the feasibility of deepGPS by focusing on single-localized proteins only in the cytoplasm (182 unique protein sequences with 8,289 cropped images in training set and 40 unique protein sequences with 1,919 cropped images in test set) or nucleoplasm (109 unique protein sequences with 5,174 cropped images in training set and 23 unique protein sequences with 1,127 cropped images in test set). This subset was used to train and evaluate the deepGPS-single-2 model (Fig. 2c and Extended Data Fig. 3), which is designed to generate both the protein localization image and the binary-classification text label. As shown in Figure. 2d, deepGPS-single-2 provided accurate protein subcellular localization predictions and generated fluorescence images (“Protein-Predicted”) that were highly similar to the ground truth fluorescence images (“Protein-GT”). Additionally, we conducted a cross-assay to evaluate the robustness of deepGPS-single-2 (Fig. 2e, left). In this assay, we used the ground-truth nucleus image from argonaute RISC component 1 (AGO1) as a nuclear fiducial marker and input the protein sequences of AGO1 (in cytoplasm), family with sequence similarity 120 member A (FAM120A, in cytoplasm), heterogeneous nuclear ribonucleoprotein D (HNRNPD, in nucleoplasm) and SWI/SNF related BAF chromatin remodeling complex subunit D2 (SMARCD2, in nucleoplasm) into deepGPS-single-2, respectively. Interestingly, each prediction closely matched the expected localization, suggesting that deepGPS-single-2 has learned the intrinsic protein localization patterns from protein sequences but independent of the nucleus image. Similar results were observed using ground-truth nucleus images from other proteins (Fig. 2e). These results indicate the excellent performance and robustness of deepGPS-single-2 in protein localization prediction.

### Evaluation of model performance by adding structure information to deepGPS

In addition to protein sequences, we tempted to supplement protein structures information into the model training and tested whether it would improve the model’s performance. Specifically, we downloaded the PDB files containing protein structure information predicted by AlphaFold2 ^16^ and converted each of them to a point cloud in tensor format using PyUUL ^18^, with three channels representing carbon, oxygen, and nitrogen atoms (Fig. 3a). The point cloud tensor was then transformed into a structure latent vector through the PointNet ^19^ and combined to generate the model output (Extended Data Fig. 4a, bottom).

**Fig. 3:**
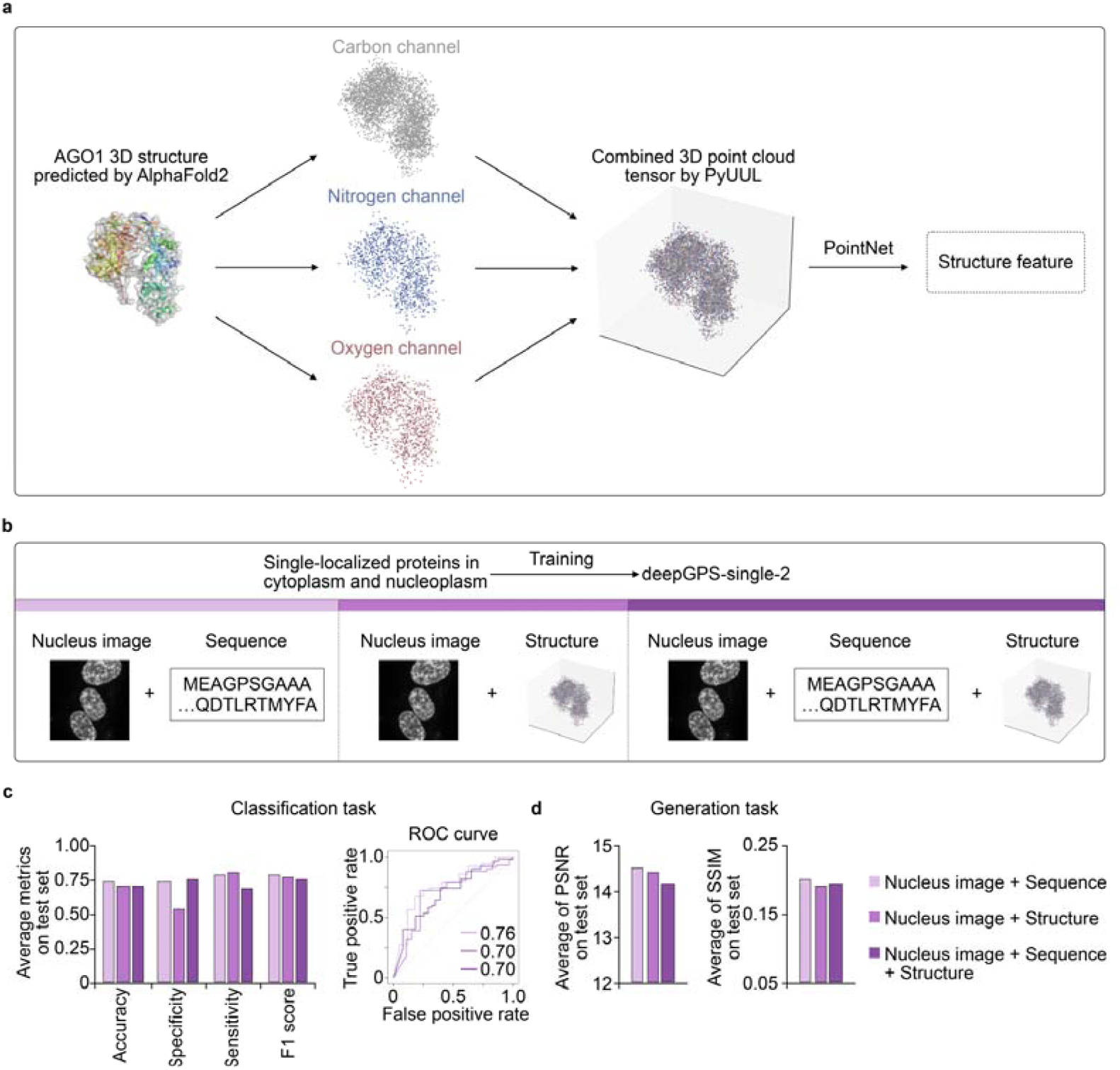
Performances of deepGPS-single-2 variants with different input formats. **a**, Schematic diagram illustrating the conversion of a protein structure predicted by AlphaFold2 into a point-cloud tensor with carbon, nitrogen, and oxygen channels using PyUUL. **b**, Strategies for training three variants of deepGPS-single-2 with different inputs of nucleus image + protein sequence, nucleus image + protein structure and nucleus image + protein sequence + protein structure. **c,** General performances of deepGPS-single-2 variants on the classification task including accuracy, specificity, sensitivity and F1 score in left and ROC curve in right. **d,** General performances of deepGPS-single-2 variants on the generation task.

Next, we compared three variants of deepGPS-single-2 with different input combinations of nucleus image + protein sequence (Fig. 3b, left and Fig. 1), nucleus image + protein structure (Fig. 3b, middle and Extended Data Fig. 4a) and nucleus image + protein sequence + protein structure (Fig. 3b, right and Extended Data Fig. 4b). Interestingly, deepGPS-single-2 with nucleus image + protein sequence as the input demonstrated the best performance in the classification task with the highest accuracy of 0.74 (Fig. 2c, left) and AUROC of 0.76 (Fig. 3c, right), and also exhibited the best performance in the image generation task with an average peak signal-to-noise ratio (PSNR) of 14.5 and an average structural similarity index measure (SSIM) of 0.20 (Fig. 3d). Interestingly, the addition of structural information from AlphaFold2 into the model training showed little impact on the improvement of the model’s performance (Fig. 3c,d). We speculated that since the protein sequence itself was used for protein structure prediction by AlphaFold2, the pre-trained large language model of protein, ESM-2, might be sufficient in extracting intrinsic features from the protein sequence for protein subcellular localization prediction. Therefore, we selected to use the nucleus image + protein sequence combination as the input for the model development.

We further compared deepGPS-single-2 with other reported protein localization prediction tools, including MultiLoc2-LowRes ^10^ and DeepLoc ^11^ in the classification task. Given that these tools output different numbers of classes, we summarized the outputs from MultiLoc2-LowRes and DeepLoc into two major subcellular localizations: cytoplasm and nucleoplasm (see the “Methods” section) for comparison. As shown in Supplementary Table S1, deepGPS-single-2 exhibited comparable performance to the other tools in the test dataset. More specifically, the AUROC of deepGPS-single-2, MultiLoc2-LowRes and DeepLoc is 0.76, 0.77 and 0.80, respectively, and the AUPRC of deepGPS-single-2, MultiLoc2-LowRes and DeepLoc is 0.84, 0.80 and 0.82, respectively. However, given that deepGPS-single-2 was trained only on a small-sized dataset with 291 protein sequences in total, far fewer than those used in MultiLoc2-LowRes and DeepLoc (4,286 and 13,858 protein sequences, respectively), the performance of deepGPS-single-2 in protein subcellular localization was robust. Strikingly, the deepGPS model can *de novo* produce protein subcellular localization images from the protein sequence in addition to textual labels. Thus, the deepGPS model is unique in protein subcellular localization prediction with generative images as an extra layer of outputs.

Together, we constructed a multimodal model, deepGPS-single-2, which could achieve the accurate protein localization prediction with both text and image outputs.

### Extended deepGPS models to predict other types of subcellular localization

Inspired by the performance of deepGPS-single-2 in predicting cytoplasmic and nucleoplasmic proteins, we further constructed deepGPS-single-4 to extend the predicting classes to include cytoplasm (182 unique protein sequences with 8,289 cropped images in training set and 40 unique protein sequences with 1,919 cropped images in test set), nucleoplasm (109 unique protein sequences with 5,174 cropped images in training set and 23 unique protein sequences with 1,127 cropped images in test set), vesicles (112 unique protein sequences with 5,126 cropped images in training set and 28 unique protein sequences with 1,326 cropped images in test set), and endoplasmic reticulum (ER) (64 unique protein sequences with 3,334 cropped images in training set and 14 unique protein sequences with 593 cropped images in test set) (Fig. 4a, top and Extended Data Fig. 3). Proteins in these classes are predominant in the OpenCell database (Fig. 2b). After training with the same strategy for deepGPS-single-2, deepGPS-single-4 also showed good performance in the classification task, with an average accuracy of 0.81 and (Fig. 4b) an average AUROC of 0.83 (Fig. 4c). Since vesicles and ER belong to the cytoplasm in biology, classification mistakes for vesicles and ER mostly occurred within these three internal classes, but were rarely classified as nucleoplasm (Fig. 4d,e). As deepGPS-single-2, deepGPS-single-4 can generate protein images with general shapes highly similar to the ground truth under different image quality cutoffs (Fig. 4f and Extended Data Fig. 5a,b), suggesting its capability for higher resolution protein localization prediction.

**Fig. 4:**
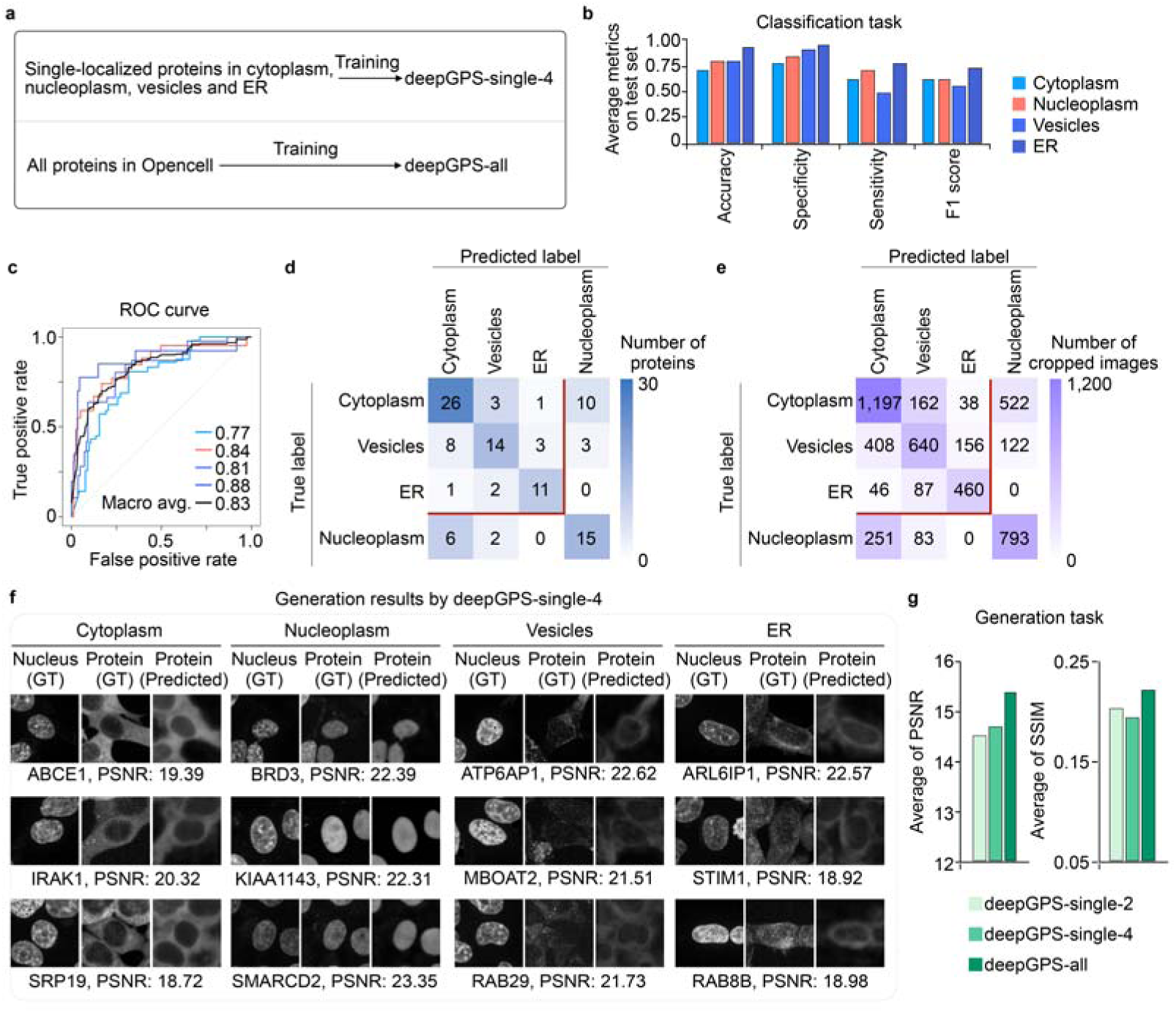
Extended deepGPS models for predicting other subcellualar localization types. **a**, Strategies for training deepGPS-single-4 and deepGPS-all. **b,c,** General performance of deepGPS-single-4 on the classification task including accuracy, specificity, sensitivity and F1 score in panel **b** and ROC curve in panel **c**. **d,e,** Confusion matrix of proteins (**d**) and cropped images (**e**) for the classification task achieved by deepGPS-single-4. **f,** Twelve instances of generation results by deepGPS-single-4. GT, ground truth; PSNR, peak signal-to-noise ratio. **g,** General performances of deepGPS-single-2, deepGPS-single-4 and deepGPS-all on the generation task.

Lastly, we set to apply all proteins and their images (1,051 unique protein sequences with 49,971 cropped images in training set and 250 unique protein sequences with 12,137 cropped images in test set), including single-localized proteins in 17 organelles and all multi-localized proteins, to train and evaluate deepGPS-all (Fig. 4a, bottom and Extended Data Fig. 3). Due to the limited number of single-localized proteins in certain hardly-discriminated organelles (such as mitochondria with 10 protein sequences, and cytoskeleton with 11 protein sequences) (Fig. 2b and Extended Data Fig. 2c) and multi-localized proteins (Extended Data Fig. 2b), we adjusted deepGPS-all for the image generation task only, not for classification. As shown in Extended Data Fig. 6, deepGPS-all can generate highly similar images in general shapes compared to the ground truth for most organelles. These included the cytoplasm, vesicles, ER, Golgi apparatus, cytoskeleton, and mitochondria, which generally showed cytoplasmic preference in biology (Extended Data Fig. 6a), as well as the nucleoplasm, chromatin, nuclear membrane, nuclear GC, nuclear FC/DFC, and nuclear punctae, which generally showed nuclear preference in biology (Extended Data Fig. 6b). These results together indicated the strong performance of deepGPS-all in protein localization with higher resolution. Of special note, comparing to deepGPS-single-2 and deepGPS-single-4, deepGPS-all showed the best image generation ability with the highest PSNR and SSIM (Fig. 4g), suggesting that deepGPS performance can be further improved with larger datasets.

So far, by constructing deepGPS-single-4 and deepGPS-all, we achieved accurate protein localization prediction with the higher resolution in organelles. However, due to the insufficient information of proteins in hardly-discriminated organelles, we expected that including additional data for the model training will further improve the model performance.

### An interactive and public online platform for protein localization prediction

To provide an open environment for protein localization research, we established the openGPS website (https://bits.fudan.edu.cn/opengps) based on deepGPS models (Fig. 5a) with Flask framework. The openGPS website contains two major function modules: “Prediction” and “Submission” (Fig. 5b). On one hand, the “Prediction” module allows online users to input the sequence of a query protein and a nucleus image (optional; a default nucleus image will be used if not available). Then, deepGPS will return the localization prediction with both a text label and an image according to the given nucleus image (or the default nucleus image), based on the selected model (deepGPS-single-2, deepGPS-single-4, or deepGPS-all) (Fig. 5c). On the other hand, the “Submission” module allows users to submit their own experimental image of protein localization to openGPS (Fig. 5d), which not only enables users to compare their results with those generated by openGPS for interest but also further expands the data volume of proteins in nuanced organelles. For convenience and efficiency, both “Prediction” and “Submission” modules support batch processing.

**Fig. 5:**
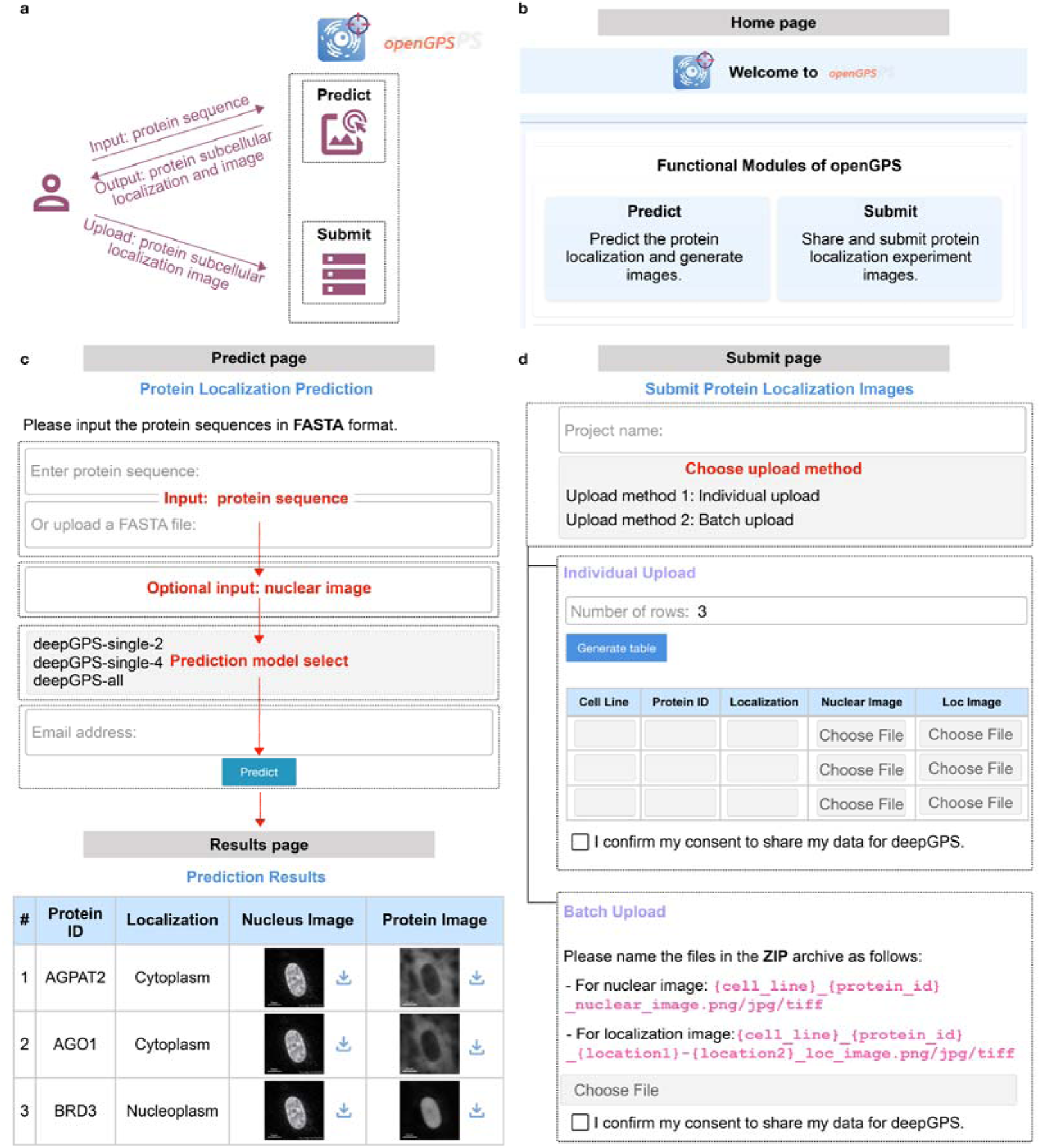
A shared platform for protein subcellular localization prediction provided by openGPS. **a**, Schematic diagram illustrating the user interaction with the openGPS website, including two functional modules of “Predict” and “Submit”. **b**, “Home” page of openGPS website. **c**, “Predict” and “Results” pages of openGPS website. After entering protein sequences and selecting the prediction model, users can submit the prediction process to receive tabulated results, which are available for download. **d**, “Submit” page of openGPS website. Users can submit true images of protein localization with “Individual Upload” or “Batch Upload” modes.

Here, we tested the “Predict” module of openGPS for the localization of additional independent proteins not included in the OpenCell database, including nucleoplasmic protein cleavage and polyadenylation specific factor 6 (CPSF6), nucleoplasmic protein cleavage stimulation factor subunit 2 (CSTF2) and cytoplasmic protein eukaryotic translation elongation factor 2 (EEF2), from RBP Image database ^20^. Despite these three proteins being absent from our training set and all originating from HeLa cell imaging, their localizations were accurately predicted in both text labels and fluorescence images (Extended Data Fig. 7), underscoring the strong generalizability of openGPS.

Together, openGPS will not only provide an effective tool for protein localization prediction but also expand the scale of datasets to refine our model, thus forming a positive feedback loop for protein functional studies.

## Discussion

Protein subcellular localization is highly associated with proteins’ function and the overall cellular dynamics ^21^. Knowing where proteins are localized will help in understanding their cellular functions and even developing medical and biotechnological applications (such as NLS for CRISPR-Cas system). With the rapid development of deep learning, a series of computational models have been developed to predict protein subcellular localization based on the primary sequence of protein^10,11^, which generally report protein subcellular localization prediction with text labels as outputs. Although fluorescence image analyses have been widely used for protein subcellular localization analyses, no text-to-image prediction model has been developed.

In this study, we designed deepGPS, a deep generative model for *de novo* <u>p</u>rotein <u>s</u>ubcellular localization prediction that uses protein sequences and nuclear fiducial markers as inputs to generate both text labels and artificial images of protein subcellular localizations as outputs (Fig. 1). By constructing a comprehensive image dataset from the OpenCell database ^12^ (Fig. 2a,b and Extended Data Figs. 2 and 3), we first trained the deepGPS-single-2 model to predict the localization of cytoplasmic and nucleoplasmic proteins (Fig. 2c). Remarkably, deepGPS-single-2 exhibited reliable performance in both image generation and text classification tasks (Figs. 2d,e and 3c,d). In addition, deepGPS-single-4 and deepGPS-all models that predict four or all types of protein subcellular localization also achieved reasonable results (Fig. 4 and Extended Data Figs. 5 and 6). Finally, we presented the openGPS website, which allows users to predict or compare their experimental results online (Fig. 5). Using the “Predict” module of openGPS, we successfully predicted the localization of three proteins from the independent experiment in HeLa cells ^20^, not included in the OpenCell database, with accurately generating both text labels and fluorescence images (Extended Data Fig. 7), highlighting the strong generalizability of our deepGPS model.

Compared to existing models for protein localization prediction, our model offers several distinct advantages: 1) Multimodal capability, whereas traditional protein localization prediction methods are text-to-text; 2) Different to text labels, images can more clearly depict the distribution of proteins within cells; 3) Image outputs can better illustrate the multi-localized proteins and are expected to provide quantitative insights into protein localization.

It should be noted that our model is currently just a prototype of a multimodal model for *de novo* predicting protein localization. Following questions need to be further addressed. Firstly, protein structure information may also play an important role in protein subcellular localization. In this study, we used the structure information from AlphaFold2 and converted it to a point cloud tensor (Fig. 3a and Extended Data Fig. 4), but additional structure information did not improve the model performance (Fig. 3c,d). Experimental structure data and appropriate encoding methods need to be tested in the future. Secondly, we observed that deepGPS currently generates images highly similar to the ground truth in general shapes. However, it is challenging to depict the texture and details of proteins in more hardly-discriminated organelles. Thus, sufficient protein data is needed to further improve the generation performance of our model. Thirdly, since the original data from OpenCell comprised proteins only in HEK293T cells, models enabling protein localization prediction across different cell lines or biological conditions are warranted.

Together, we present a generative model for *de novo* predicting protein localization based on a multimodal deep learning model, and further provide a public and convenient online website, openGPS, for the protein localization research community. It is expected that expanding the scale of datasets will further refine the model for better prediction outcomes.

## Methods

### Fluoresence image dataset of protein localization prediction

All fluorescence images of proteins were downloaded from OpenCell ^12^, which encompasses 6,301 nucleus-protein paired 2D images (600 × 600 pixels) for 1,311 tagged proteins. Given that AlphaFold2 provides predicted protein structures with multiple overlapping fragments when the input protein exceeds 2,700 amino acids, we selected only proteins shorter than 2,700 amino acids, which includes 6,239 paired images of 1,301 proteins, to construct the image dataset for protein localization prediction (Extended Data Fig. 3).

### Image data preprocessing

To improve the quality and expand the size of the image dataset, the raw pixel intensities of the images were first normalized using CSBDeep ^22^. Then, nuclei in the normalized images were segmented using StarDist^17^, resulting in all pixels belonging to one nucleus region being labeled as a specific group. Some noise dots that were incorrectly segmented as nuclei were removed through size selection if their labeled regions were smaller than 1,000 pixels. All denoised images were cropped to 208 × 208 pixels, ensuring at least one nucleus was centered in the image. Finally, a total of 62,108 cropped images of nuclei and paired proteins were obtained for subsequent model construction and evaluation.

### Annotation of protein localization

For the text label of protein localization, we utilized manually assigned subcellular localization categories for each tagged protein from the OpenCell library^12^. In this annotation, each category is divided into three grades: grade 3 indicates very prominent localization, grade 2 represents unambiguous but less prominent localization, and grade 1 denotes weak or barely detectable localization. We considered the localization recorded in the highest available grade (grade 3; if unavailable, grade 2) as the major localization of a given protein. Of note, in this study, a protein with a single major localization was defined as a single-localized protein, while a protein with multiple major localizations was defined as a multi-localized protein

### Dataset split for model training and evaluation

To train the deepGPS-single-2 and deepGPS-single-4 models, we used only cropped images of single-localized proteins. Since the number of proteins varied across localization classes, we first sorted all classes by the number of proteins they contained. Then, using a linear sampling method, we randomly selected approximately 75% to 90% of the proteins from each class as the training set. The remaining proteins were evenly split to create the validation and test sets. For the deepGPS-single-2 model, we further selected proteins localized in the nucleoplasm or cytoplasm, while for the deepGPS-single-4 model, proteins localized in the nucleoplasm, cytoplasm, ER and vesicles were selected for training and evaluation.

To train the deepGPS-all model, we used cropped images of all proteins with single or multiple major localizations. The entire dataset was randomly split into training and test sets in a 4:1 ratio, ensuring that each localization class had a sufficient number of proteins in the test set for model evaluation.

Notably, to ensure the generality and robustness of the model, the proteins in the training and test sets were unique to each group, with no overlap.

### Feature representation of the protein primary sequence

We utilized the pre-trained protein language model, ESM2-650M^14^, to capture intrinsic information from protein primary sequences. Briefly, each protein sequence was first encoded as one-hot tensor and subsequently converted into a vector of dimensions *L* × 1280 by ESM2-650M, where *L* represents the number of amino acids in a given protein sequence. Given the variable length of protein sequences and the fact that the mean of the vector from ESM2 can serve as an overall representation of the sequences^23^, each protein was finally represented by a vector *f* ∈ *R*^1280^ by reducing the dimensionality using mean pooling. To ensure the extracted features are better suited to our task, we apply a linear layer to transform the features into *f_s_* ∈ *R*^1024^.

### Feature representation of the protein tertiary structure

To integrate the information of protein tertiary structure in our model, we first downloaded the predicted structures (PDB files) of proteins in our dataset from AlphaFold2 database ^24^. Each PDB file was then converted to an *n* × 6 point cloud tensor using PyUUL^18^, where n is the number of points. The first three of the six dimensions represent the 3D coordinates of each point, while the last three dimensions represent the carbon, oxygen and nitrogen atoms, respectively. Finally, the point cloud tensor was converted to a global representation *f_p_* ∈ *R*^1024^ by using PointNet output ^19^, capturing the global geometric and structural properties of the input protein.

### Image generation by deepGPS

U-Net^25^ was used as the foundational framework for the generation task in the deepGPS model. Specifically, the U-Net network comprises a down-sampling module (encoder) and an up-sampling module (decoder). First, the encoder extracts features from the input image while reducing its spatial dimensions, consisting of four convolutional blocks. Each block contains two convolutional layers followed by a max-pooling layer. This process yields the feature 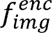 ∈ *R*^1024×13×13^.

Of note, if the input consists solely of protein sequence feature *f_s_*, the latent vector *f_hidden_* is represented by *f_s_*. If the input includes both a 3D protein structure representation *f_p_* and a sequence feature *f_s_*, *f_p_* and *f_s_* are concatenated to generate new hidden features *f_hidden_* ∈ *R*^2×1024^.

Next, to facilitate fusion with the latent vector *f_hidden_*, 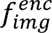 is reshaped into *f_enc_* ∈ *R*^1024×169^. The fusion process employs an attention mechanism where *f_enc_* serves as the Query (Q), and *f_hidden_* acts as both the Key (K) and Value (V). The resulting fused feature is *f_fuse_* ∈ *R*^1024×169^, computed as:

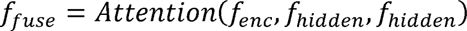

The decoder then reconstructs the image from *f_fuse_*. Each step in the decoder involves up-sampling the features, applying convolution to halve the number of feature channels, and concatenating the result with the corresponding feature map from the encoder. After performing two 3 × 3 convolutions, each followed by a ReLU activation, an image of the same size as the input is finally generated.

## Data availability

The images and annotations of protein localization used in this work are available from the OpenCell database (https://opencell.czbiohub.org/download). Predicted 3D protein structures are available from the AlphaFold database (https://alphafold.ebi.ac.uk/download).

## Code availability

All codes for constructing deepGPS models has been deposited at Github at https://github.com/royal-dargon/deepGPS/.

## Acknowledgements

We thank all lab members for their critical discussion. This work was supported by grants from the National Natural Science Foundation of China (NSFC, 31925011 to L.Y.), and the Science and Technology Commission of Shanghai Municipality (STCSM, 23JS1400300 and 23DX1900102 to L.Y.). This work was also partially supported by Shanghai Artificial Intelligence Laboratory.

## Contributions

L.Y. and G.-H.Y. conceived and designed the project. L.Y. and N.D. supervised the project; L.Y. managed the project; J.L. constructed the deepGPS model and performed model evaluation with the help of Z.Yang and Z.Yuan, supervised by N.D.; G.-H.Y. analysed the data, supervised by L.Y.; Y.-Q.C. constructed openGPS website with the help of G.-H. Y., supervised by L.Y.; W.O. and T.C. provided support with techniques; G.-H.Y. and L.Y. prepared figures and wrote the paper with input from all authors. All authors read and approved the final manuscript.

## Ethics declarations

Competing interests

The authors declare no competing interests.

## EXTENDED DATA

**Extended Data Fig. 1:**
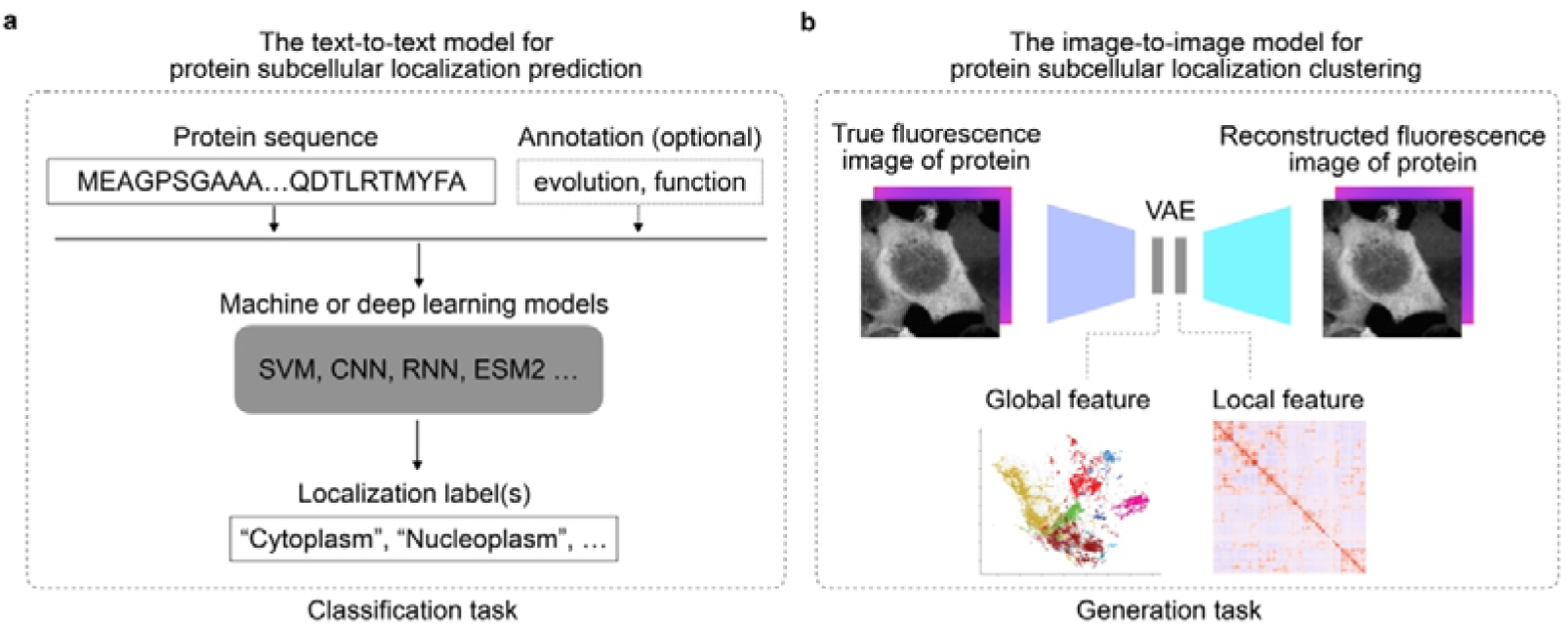
Other text-to-text and image-to-image models for protein subcellular localization prediction or clustering. **a**, A schematic diagram of the text-to-text model for predict protein subcellular localization prediction. **b**, A schematic diagram of the image-to-image model for protein subcellular localization clustering, referred to Cho *et al*. (PMID: 35879608).

**Extended Data Fig. 2:**
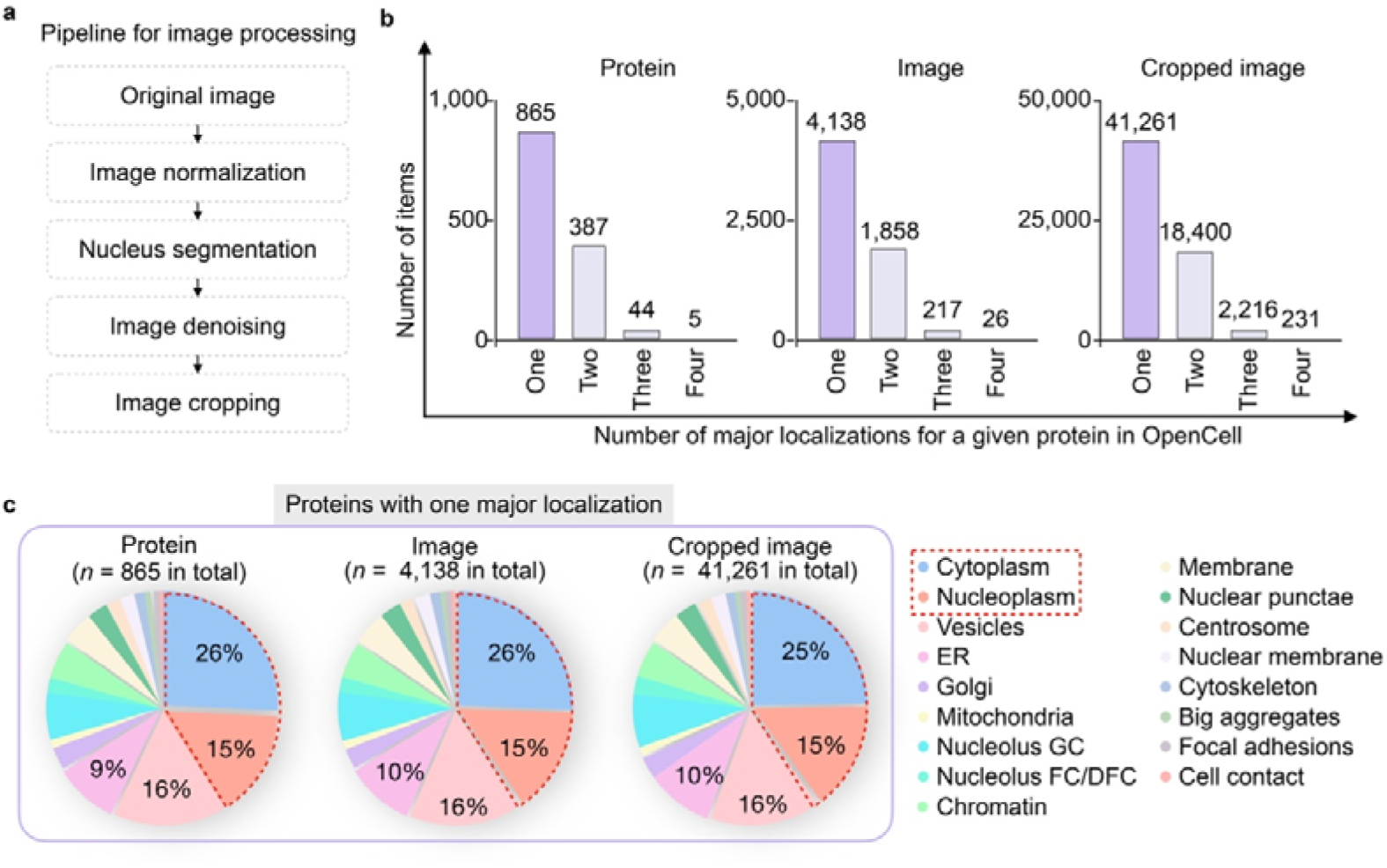
Construction of image-based protein localization dataset. **a**, Pipeline for image processing. **b**, Numbers of proteins (left), original images (middle) and cropped images (right) with different major localizations. **c**, Distribution of proteins, original images (middle) and cropped images (right) with only one major localization.

**Extended Data Fig. 3.**
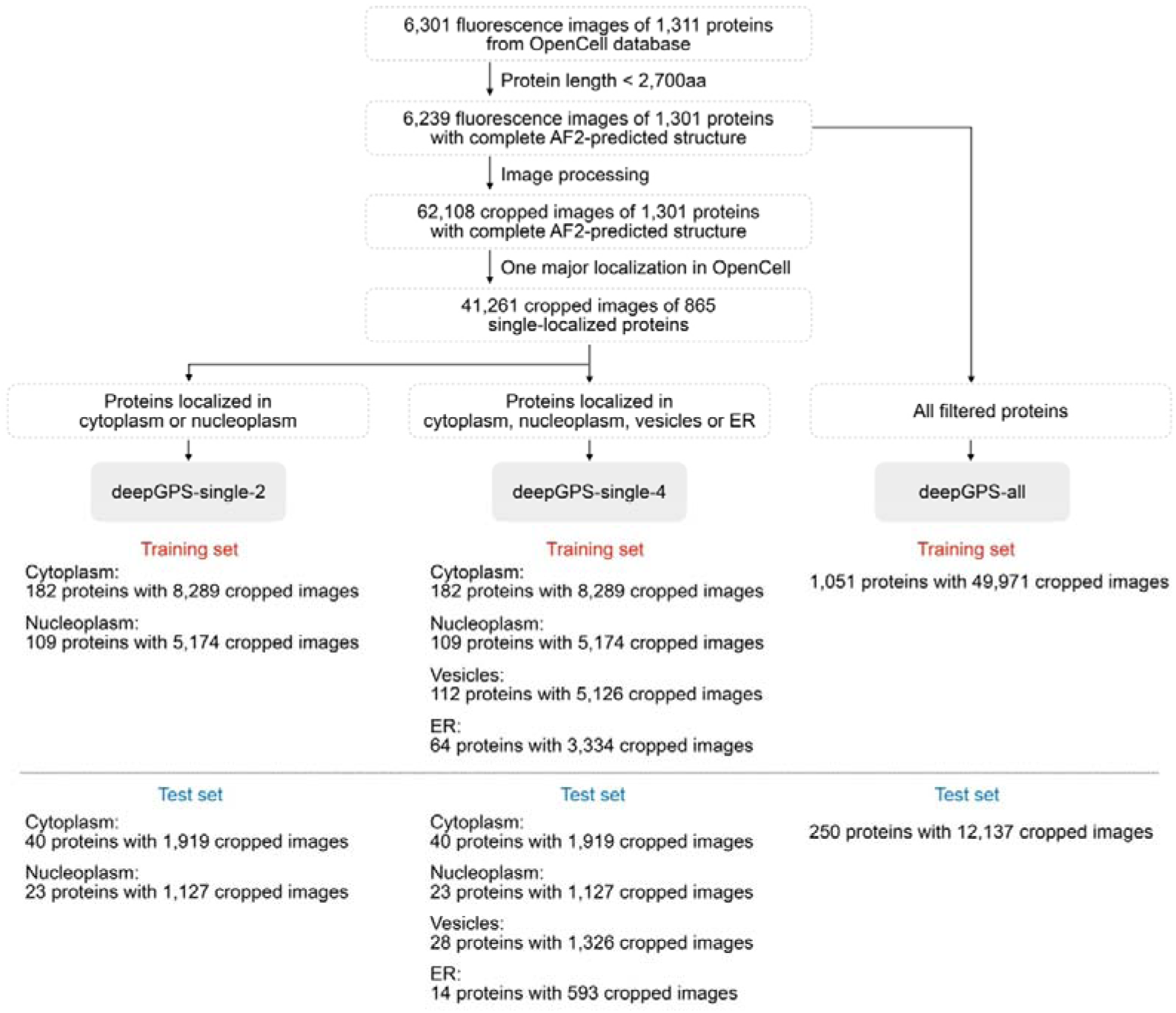
Datasets for training and evaluating deepGPS models. Pipeline of constructing datasets used for training and evaluating deepGPS-single-2, deepGPS-single-4 and deepGPS-all models.

**Extended Data Fig. 4:**
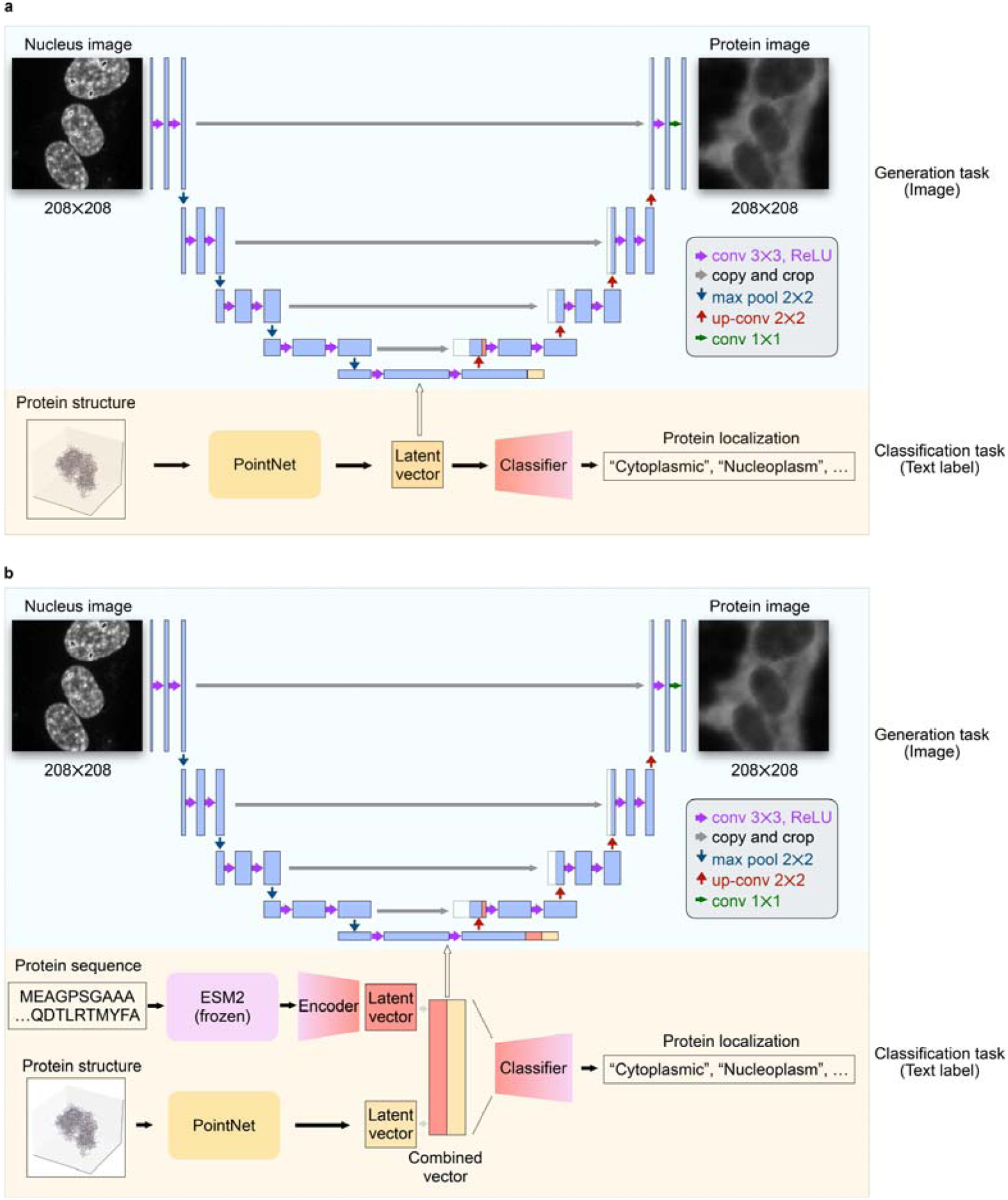
Training deepGPS with additional information of protein 3D structure. **a**, Schematic diagram illustrating the architecture of deepGPS with the nucleus image and protein structure as inputs. **b**, Schematic diagram illustrating the architecture of deepGPS with the nucleus image, protein sequence and protein structure as inputs.

**Extended Data Fig. 5:**
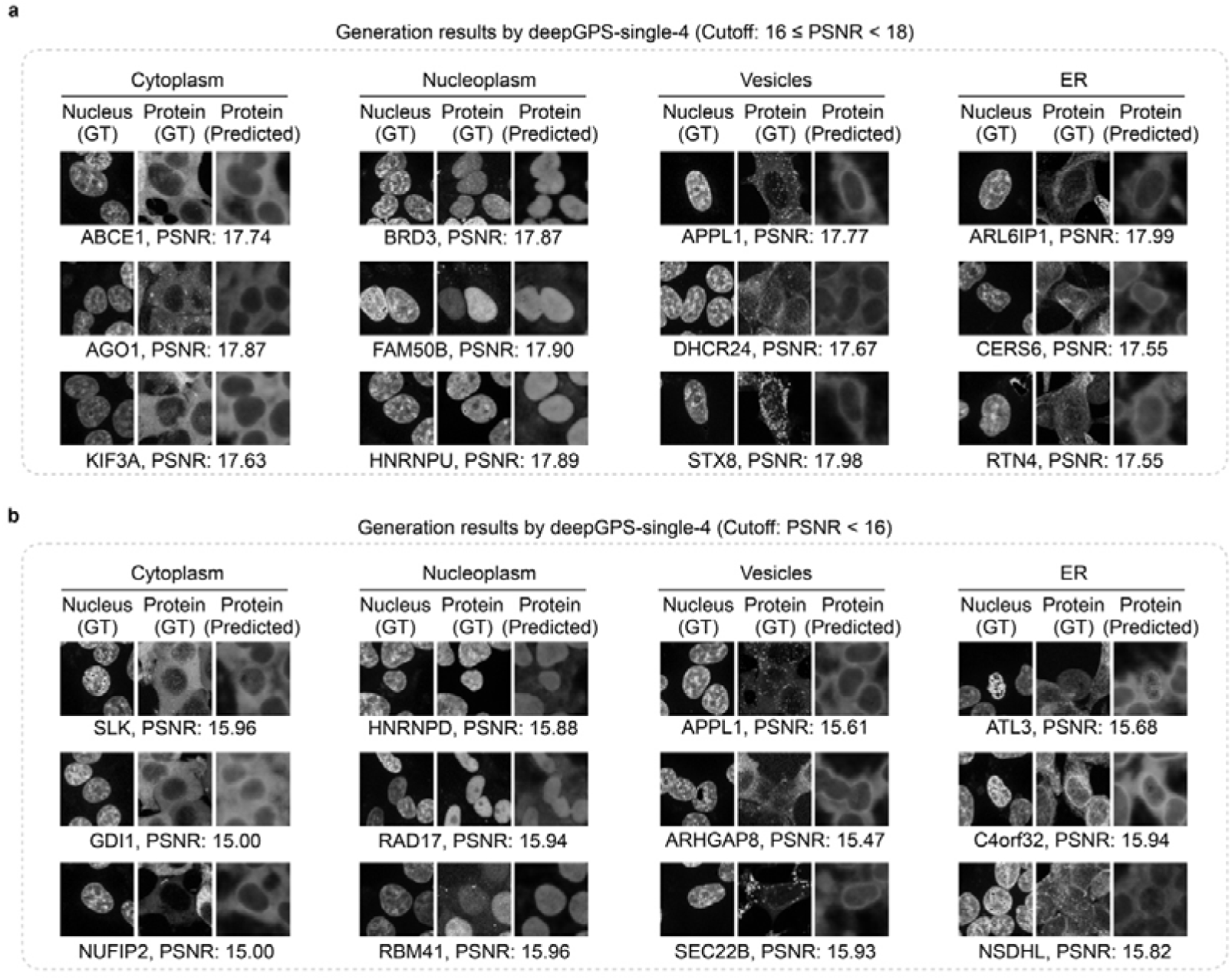
**Performance of deepGPS-single-4 on predicting protein subcellular localization. a,b**, Specific image instances generated by deepGPS-single-4 under the different image qualities of 16 ≤ PSNR ≤ 18 (**a**) and PSNR < 16 (**b**). Of note, the proteins corresponding to images generated by deepGPS in panels **a** and **b** were all from test set, with sequences that were not used during deepGPS training.

**Extended Data Fig. 6:**
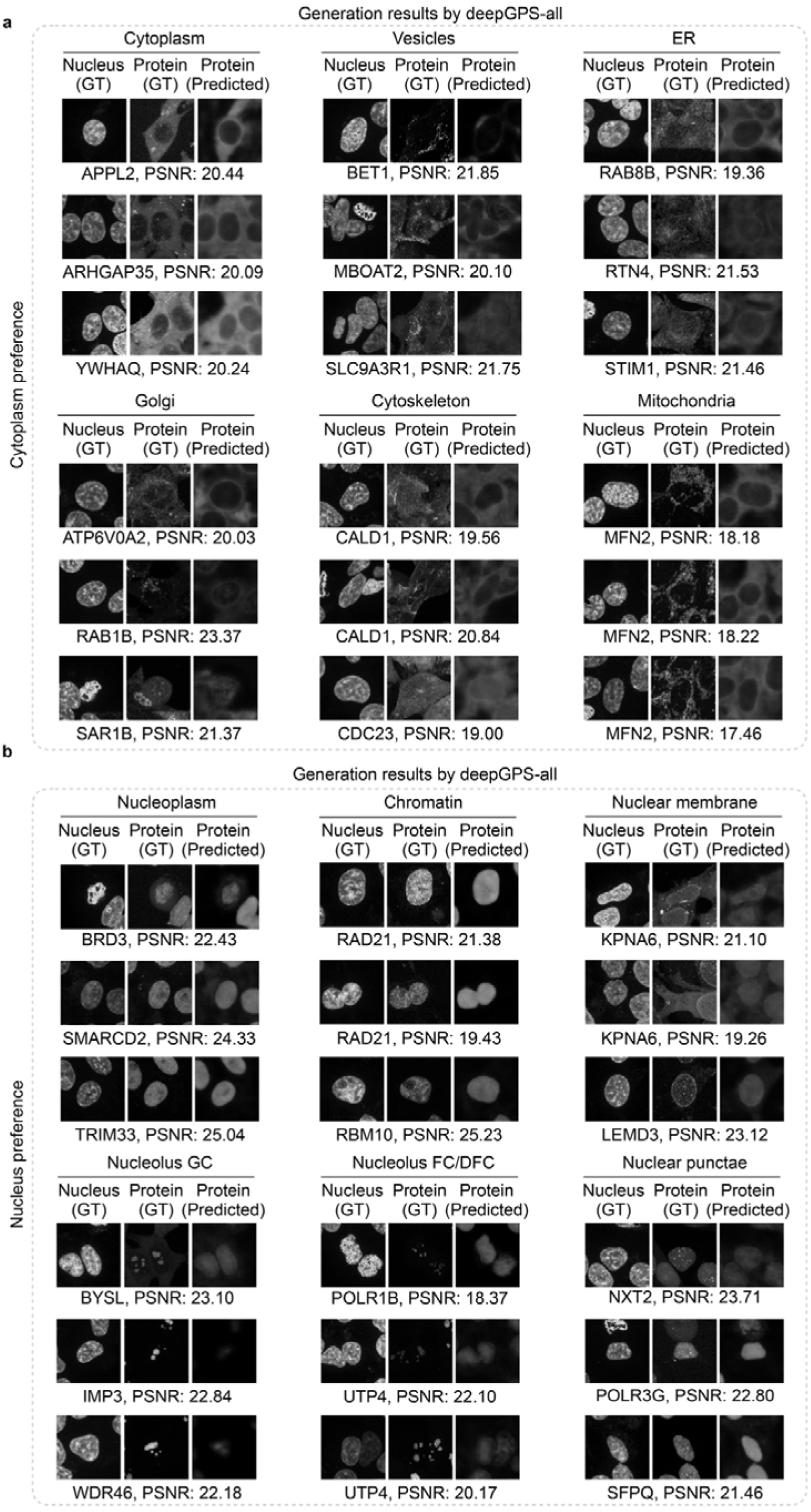
Protein localization images generated by deepGPS-all. **a**, Protein localization images generated by deepGPS-all, showing a preference for proteins primarily localized in the cytoplasm. **b**, Protein localization images generated by deepGPS-all, showing a preference for proteins primarily localized in the nucleus. Of note, the proteins corresponding to images generated by deepGPS in panels **a** and **b** were all from test set, with sequences that were not used during deepGPS training.

**Extended Data Fig. 7:**
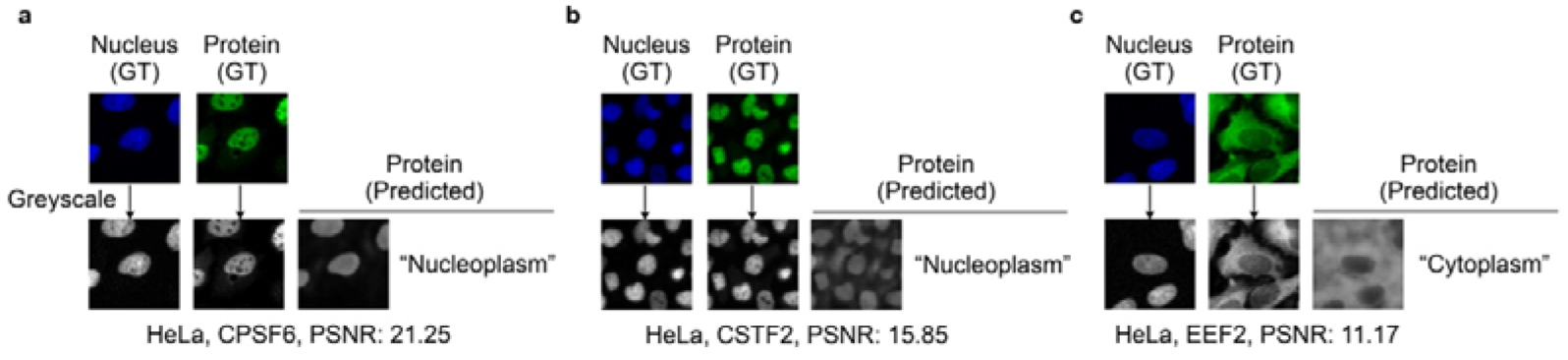
Prediction for independent proteins with openGPS. a-c, Predicted results of three independent proteins from RBP Image Database, including CPSF6 (**a**), CSTF2 (**b**) and EEF2(**c**). GT, ground truth; PSNR, peak signal-to-noise ratio.

## References

1. Tokuriki, N. & Tawfik, D.S. Protein dynamism and evolvability. Science 324, 203–207 (2009).

2. Liu, Y., Beyer, A. & Aebersold, R. On the Dependency of Cellular Protein Levels on mRNA Abundance. Cell 165, 535–550 (2016).

3. Hetz, C., Zhang, K. & Kaufman, R.J. Mechanisms, regulation and functions of the unfolded protein response. Nat Rev Mol Cell Biol 21, 421–438 (2020).

4. Sellars, M.C., Wu, C.J. & Fritsch, E.F. Cancer vaccines: Building a bridge over troubled waters. Cell 185, 2770–2788 (2022).

5. Kowall, N.W. & Kosik, K.S. Axonal disruption and aberrant localization of tau protein characterize the neuropil pathology of Alzheimer’s disease. Ann Neurol 22, 639–643 (1987).

6. Jiao, W. et al. Aberrant nucleocytoplasmic localization of the retinoblastoma tumor suppressor protein in human cancer correlates with moderate/poor tumor differentiation. Oncogene 27, 3156–3164 (2008).

7. Cao, S.M. et al. Altered nucleocytoplasmic export of adenosine-rich circRNAs by PABPC1 contributes to neuronal function. Mol Cell 84, 2304–2319 e2308 (2024).

8. UniProt, C. UniProt: the Universal Protein Knowledgebase in 2023. Nucleic Acids Res 51, D523–D531 (2023).

9. Hua, S. & Sun, Z. Support vector machine approach for protein subcellular localization prediction. Bioinformatics 17, 721–728 (2001).

10. Blum, T., Briesemeister, S. & Kohlbacher, O. MultiLoc2: integrating phylogeny and Gene Ontology terms improves subcellular protein localization prediction. BMC Bioinformatics 10, 274 (2009).

11. Almagro Armenteros, J.J., Sonderby, C.K., Sonderby, S.K., Nielsen, H. & Winther, O. DeepLoc: prediction of protein subcellular localization using deep learning. Bioinformatics 33, 4049 (2017).

12. Cho, N.H. et al. OpenCell: Endogenous tagging for the cartography of human cellular organization. Science 375, eabi6983 (2022).

13. Kobayashi, H., Cheveralls, K.C., Leonetti, M.D. & Royer, L.A. Self-supervised deep learning encodes high-resolution features of protein subcellular localization. Nat Methods 19, 995–1003 (2022).

14. Lin, Z. et al. Evolutionary-scale prediction of atomic-level protein structure with a language model. Science 379, 1123–1130 (2023).

15. Lin, P., Yan, Y. & Huang, S.Y. DeepHomo2.0: improved protein-protein contact prediction of homodimers by transformer-enhanced deep learning. Brief Bioinform 24 (2023).

16. Jumper, J. et al. Highly accurate protein structure prediction with AlphaFold. Nature 596, 583–589 (2021).

17. Weigert, M. & Schmidt, U. Nuclei instance segmentation and classification in histopathology images with StarDist. arXiv:2203.02284 (2022).

18. Orlando, G. et al. PyUUL provides an interface between biological structures and deep learning algorithms. Nat Commun 13, 961 (2022).

19. Qi, C.R., Su, H., Mo, K. & Guibas, L.J. PointNet: Deep Learning on Point Sets for 3D Classification and Segmentation. arXiv:1612.00593 (2016).

20. Van Nostrand, E.L. et al. A large-scale binding and functional map of human RNA-binding proteins. Nature 583, 711–719 (2020).

21. Lundberg, E. & Borner, G.H.H. Spatial proteomics: a powerful discovery tool for cell biology. Nat Rev Mol Cell Biol 20, 285–302 (2019).

22. Weigert, M. et al. Content-aware image restoration: pushing the limits of fluorescence microscopy. Nat Methods 15, 1090–1097 (2018).

23. Luo, Z. et al. Interpretable feature extraction and dimensionality reduction in ESM2 for protein localization prediction. Brief Bioinform 25 (2024).

24. Varadi, M. et al. AlphaFold Protein Structure Database in 2024: providing structure coverage for over 214 million protein sequences. Nucleic Acids Res 52, D368–D375 (2024).

25. Ronneberger, O., Fischer, P. & Brox, T. U-Net: Convolutional Networks for Biomedical Image Segmentation. arXiv:1505.04597 (2015).

